# A data engine for AI-assisted labeling of high-density sperm microscopy videos

**DOI:** 10.64898/2026.11.25.769178

**Authors:** David Hart, Jayath Premasinghe, Martin Bier, Kylie Cashwell, Cameron Schmidt

## Abstract

Computer-aided semen analysis (CASA) systems are the standard tools for quantifying sperm motility, concentration, and morphology, but they rely on classical algorithms that struggle with overlapping cells, collisions, and the imaging artifacts that accumulate in long, high-density recordings. As a result, analyses are typically restricted to short videos of diluted samples. AI-based trackers could overcome these limitations, but they require large, accurately labeled training datasets, and manually labeling crowded sperm videos is prohibitively slow. Here we present a data engine for sperm tracking, where an AI model can be trained from low-density data to assist in labeling higher-density cases. We first obtain ground-truth trajectories from easy-to-label, low-density videos using a classical tracker (Trackpy) followed by human correction. From these trajectories, we generate synthetic videos that reproduce the motility characteristics of real sperm, and we further augment the real data by compositing existing videos together. This combined dataset trains an initial YOLO-based detector paired with ByteTrack, which then serves as an assistive labeler for the denser tier of videos. Through this process, we found the trained AI model was able to eliminate many of the crossovers that prevent effective labeling of sperm trajectories, with the crossover failure rate going from 44.8% down to 6.64% on medium density videos, and from 74.0% to 46.68% on high density videos.

We release the videos, corrected trajectory datasets at each density, a purpose-built labeling tool, trained models with weights, and a software package for labeling, synthetic and augmented video generation, training, and evaluation. This framework provides a practical path toward AI-based semen analysis that remains reliable at high sperm densities and over long recordings.

**Author summary:** Doctors and researchers evaluate male fertility partly by watching sperm under a microscope and measuring how they move. Automated software does this today, but it works well only on short videos of diluted samples. In crowded, longer videos, sperm cross paths and the software loses track of individual cells. Modern artificial intelligence could handle these harder videos, but it must first learn from many videos in which every sperm’s path has been carefully labeled. Creating those labels by hand for crowded videos is extremely slow. We tackled this problem by building a generated dataset from synthetic and augmented versions of sparse, easy to label real videos. Using this dataset allowed for training an AI model that was able to label higher density videos with fewer mistakes and errors, significantly speeding up the labeling process. We are sharing the videos, labels, software, and trained models so other groups can build on them.

## Introduction

Computer-aided sperm analysis (CASA) has its origins in the mid 1950s, when capillary assays were used to visually observe the linear migration of sperm [1]. By the 1970s, film- and tape-based methods enabled semi-automated analysis [2], and in the 1980s digital computers made it possible to commercialize integrated systems that combined image acquisition with automated analysis [3]. Digital imaging and microscope technology have continued to co-evolve, and in recent years a growing number of low-cost, increasingly compact imaging platforms have been adopted for clinical and veterinary use [4–6]. CASA systems are now the standard tools for evaluating sperm motility, concentration, and morphology in both research and clinical settings [7].

Despite these advances in microscope and camera technology, the kinematic analysis of sperm motion has changed little since the 1970s. Modern kinematic measurements include curvilinear velocity (VCL), straight-line velocity (VSL), average path velocity (VAP), amplitude of lateral head displacement (ALH), beat-cross frequency (BCF), and derived quantities such as linearity (LIN) [8]. Early work also revealed that some sperm exhibit a distinct form of motility, characterized by circular trajectories and asymmetric flagellar beating [9]. This ‘hyperactivated’ motility has since become associated with fertility competence, and it is typically identified by clustering kinematic measurements [10].

CASA is widely used in research and forms the basis of sperm functional analysis. In clinical practice, however, it remains secondary to other methods, such as qualitative assessment by a trained technician, owing to limited reproducibility across andrology laboratories and a relatively weak correlation between kinematic measures and fertility outcomes [11]. This is perhaps unsurprising, because ‘motility’ as defined by CASA-derived kinematics is only one aspect of sperm function. Other important correlates may include acrosomal exocytosis and chromatin condensation [12, 13]. The widespread adoption of intracytoplasmic sperm injection (ICSI), which circumvents the need for sperm motility altogether, has further reduced reliance on motility as an indicator of fertility competence [14]. ICSI nevertheless addresses extreme cases such as absolute asthenozoospermia, whereas motility remains the most practically accessible fertility indicator for most men.

The disconnect between motility and fertility status may lie in how drastically standard kinematic analysis oversimplifies sperm behavior. A typical CASA measurement observes sperm for approximately 1–2 s [7], in which time a human sperm travels only one to three body lengths [10]. Fertilization in vivo, by contrast, requires a minimum of approximately 45 minutes, and sperm must traverse a distance of 3000 body lengths or more [15]. The conventional ‘motility’ measure therefore provides only a brief snapshot of sperm behavior and does not connect sperm movement patterns to the random search behaviors that are most relevant to the physiological condition [16–19].

Long-duration tracking of sperm remains an unsolved problem for current CASA software. These systems generally rely on classical tracking algorithms that detect cells in each frame and link detections across frames using relatively simple association rules [8]. Such algorithms struggle with common imaging problems that become more severe in long videos and at high sperm densities, including cells that cross paths or overlap, transient occlusions, cells that drift out of the focal plane, and phase contrast imaging artifacts such as debris that resemble sperm. Each of these can produce missed detections or identity switches that corrupt trajectories [20]. As a result, CASA analyses typically rely on very short observation periods, and use of heavily diluted sperm samples, removing them from their native semen context. These restrictions confine traditional CASA to conditions that are poorly matched to the long-timescale search behavior described above.

AI-based detection and tracking models offer a promising solution to these limitations [21]. We anticipate that the next generation of semen analysis software will be AI-based, will handle long timescale videos at high resolution, and will be capable of accurate trajectory reconstruction at the high sperm densities found in undiluted semen samples. However, such models will require large amounts of training data to reach these performance demands, and manual labeling of trajectories in crowded videos is extremely impractical if not impossible as it may involve curation of hundreds of sperm over thousands of frames. This creates a circular bottleneck problem where the same AI techniques that would make labeling tractable are the ones that the labeled data are needed to train.

Here, we address this bottleneck with a data engine pipeline, modeled on the approach used to construct the large, high-quality training dataset of the Segment Anything Model (SAM) [22]. In a data engine, a model trained on an initial dataset is used to assist in labeling the next, more difficult tier of data, and the newly labeled data are then used to improve the model. We begin with easy-to-label, low-density, diluted sperm videos, from which we obtain an initial set of ground-truth trajectories. From these trajectories and their corresponding videos, we generate synthetic videos with motility characteristics that mirror the kinematics observed in the real data. We also create augmented real videos by combining transformed trajectory data from existing videos. Together, the synthetic and augmented videos provide sufficient training data to train an initial AI model. This model then serves as an assistive labeler for the next tier of higher-density sperm videos, on which it performs significantly better than the baseline tracker. If desired, the newly labeled real data can be used to retrain or fine-tune the model for continued improvement. This process is illustrated in Fig. 1.

**Fig 1.**
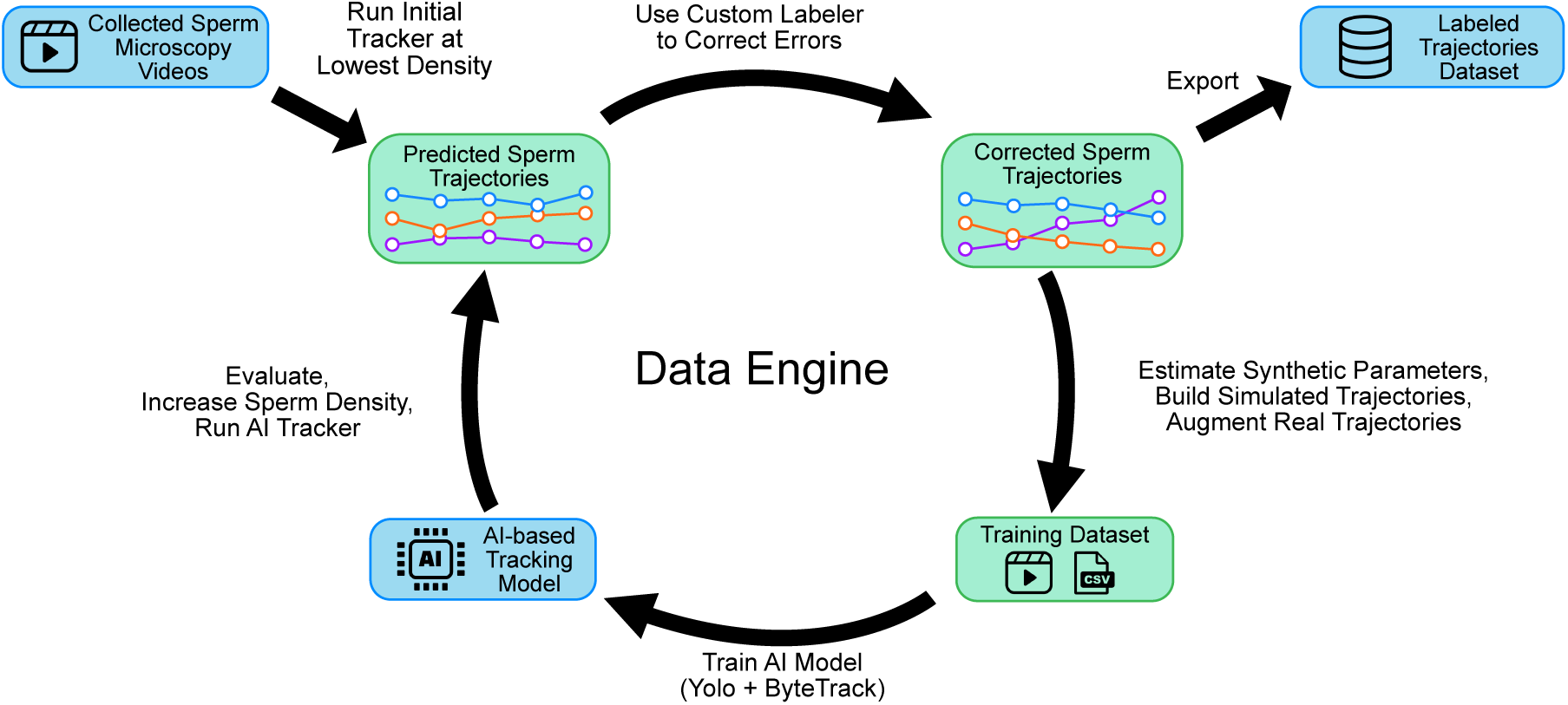
An overview of our approach. Sperm microscopy videos are collected and diluted to represent various sperm densities. Only the lowest densities can be initially tracked and corrected using classical approaches. Using synthetic and augmented generation techniques, new videos can be generated at high densities while maintaining ground truth labels. This creates a training dataset that can be used to create an AI-based tracking model. This AI model can be used to track the real data at higher densities with less cross-over errors and can be hand corrected in a fraction of the time. The boxes in blue indicate direct deliverables from this work.

## Materials and methods

### Ethics Statement

All procedures involving human subjects were approved by the Institutional Review Board of the Brody School of Medicine at East Carolina University (study ID: UMCIRB 20-001862). All samples were collected in accordance with the Declaration of Helsinki and with informed consent.

### Data Collection, Participants, and Microscopy Setup

Data was collected at East Carolina University from male volunteers ages 18-40. Semen samples were incubated for 30 minutes at 37*^◦^C*. Microscopy videos were collected on a Zeiss Axervert.A1 compound light microscope fitted with a thermal stage (Carl Zeiss AG, Jena, Germany). Full details of the data collection microscopy setup can be found in our prior work [20]. Briefly, negative phase contrast videos were obtained at 9 frames per second with 1 ms exposure on a cooled ccd array monochrome camera (Zeiss axiocam 702). The objective used was an infinity 5X magnification achromat with NA=0.24. Notably, though frame rates below 9 frames per second lack the sampling resolution for short timescale movement analysis (i.e., ALH or flagellar waveform analysis), the trajectories are sufficient for examining long timescale kinematics over periods of 30 seconds or more. Samples were loaded in a pre-warmed 20*µm* depth-chamber slide (2-XCel; Hamilton Thorne, Beverly, MA, USA). Slides were mainained during imaging on a Peltier controlled thermal stage at 37*^◦^C*. Dilutions from semen were made using human tubal fluid medium. The average loading concentrations for the various videos were: Low (1:16 dilution): 1.15 *×* 10^6^ *cells/mL*; Medium (1:8 dilution): 7.23 *×* 10^6^ *cells/mL*; and High (undiluted): 34.5 *×* 10^6^ *cells/mL*.

### Initial Sperm Trajectory Tracking

The baseline detector, tracker, and dataset were discussed previously in our work [20], but we summarize it briefly here. The baseline detecting and tracking pipeline is built on top of the Trackpy library [23]. Implementation features for filtering detections and tracks, as well as memory for disconnected tracks, were used. The parameters for detection and tracking were empirically determined given the resolution, frame rate, and speed of the sperm in our videos. Our code for our baseline tracker and data are provided at github.com/CAS-ReproLab/Sperm_Object_Tracking. For the initial dataset, five low-density videos were obtained by using a 1:16 dilution factor from collected samples. This density provided videos with only a small number of crossovers, minimizing the time needed for a human labeler to hand correct the predicted trajectories from the baseline tracker.

### Using the Custom Labeler to Correct Trajectories

A custom labeling program was built to correct trajectories from Trackpy and the improved trackers in future steps. This labeling program is built on the PyQt library [24] and has several features to simplify the labeling process for human labelers. Those features include:

- Playback and overlay controls for efficient visualization of the original data and trajectories.
- Clickable video for easy sperm ID selection.
- Built in operations for merging, splitting, swapping, and deleting sperm IDs and associated trajectories.
- A search menu for sperm by ID
- A crossover estimator function that looks for frames and sperm IDs that may have possible ID swaps based on proximity.
- Quality-of-life improvements such as hotkeys, sperm isolation, color re-randomization, and trajectory interpolation.

These features make the labeler efficient and intuitive to use. Our labeler program is visualized in 2.

### Creating Synthetic Videos

Even with a well-designed interface, manual correction of trajectories in videos is time intensive. The time required to build large datasets at the densities required is too time prohibitive. Thus, we propose two methods to increase the amount and density of the data. The first approach is to synthetically generate sperm microscopy videos with known sperm trajectories.

Building on our previous work [17, 18], we propose the following Langevin setup for generating synthetic sperm cells trajectories:

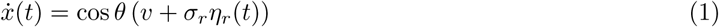

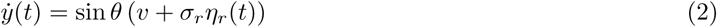

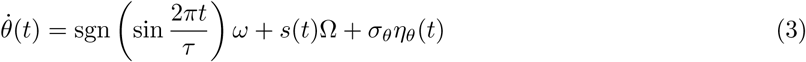

where (*x*(*t*)*, y*(*t*)) is the position of the sperm cell in the observed plane and *v* is the deterministic self-propelling velocity. The angular velocity, *θ̇*, has a short-term component, *ω*, that is due to the flagellar beating. That beating has a period *τ* . The large-scale chirality of many trajectories is described with Ω. Whether the chirality is left or right is next accounted for through *s*(*t*) = *±*1. The Gaussian noise is implemented through a normalized function *η_α_*(*t*), i.e. *< η_α_*(*t*)*η_β_*(*t^′^*) *>*= *δ_αβ_δ*(*t − t^′^*), and amplitudes *σ_r_* and *σ_θ_*. Employing Gaussian noise as the source of fluctuations means that the model is described by the standard theory of Stochastic Differential Equations [25, 26] and consistently preserves its properties when the time scale at which the process is probed is changed. A visualization of the model parameters and possible movement behaviors that can be simulated are shown in Figure 3.

**Fig 2.**
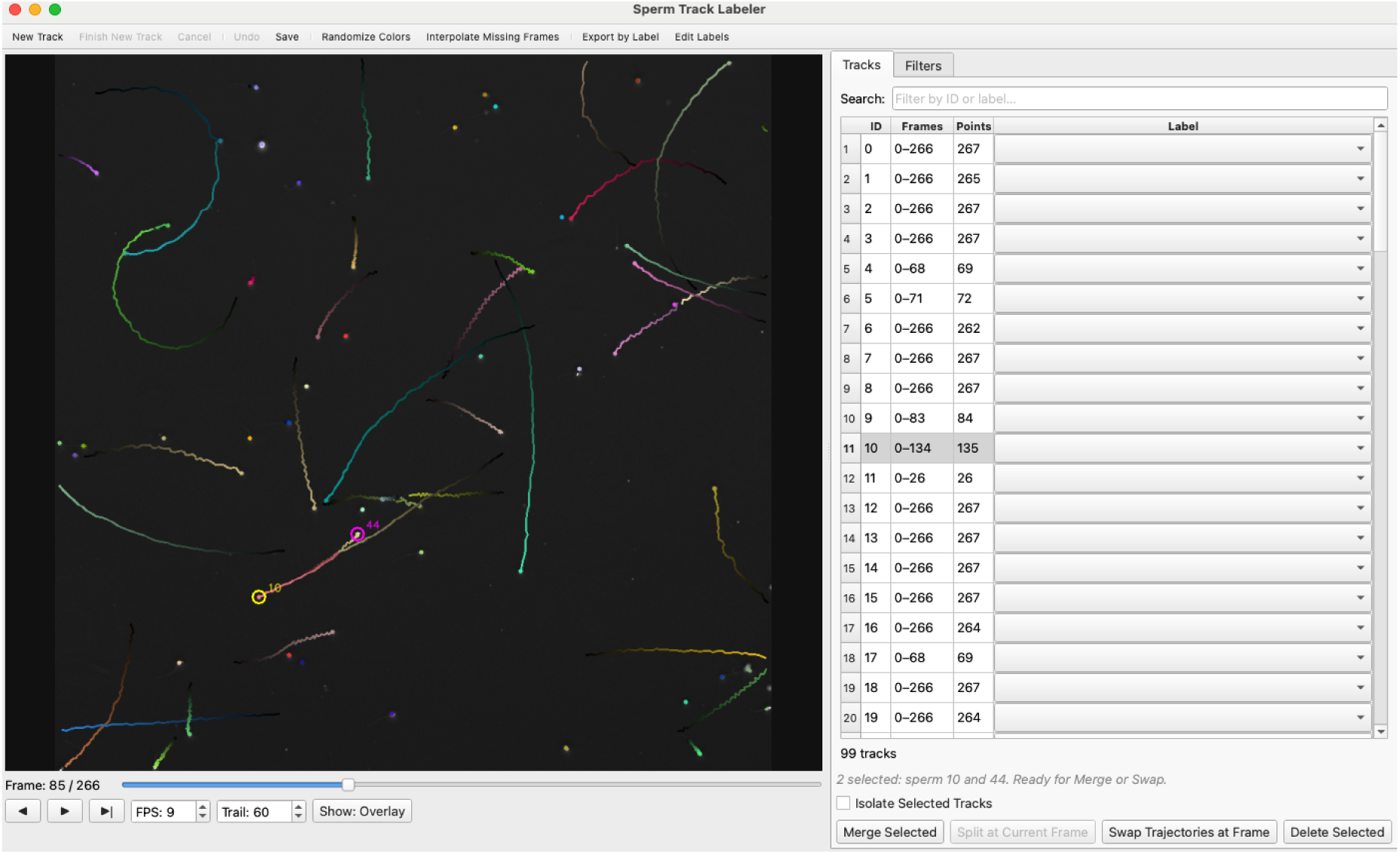
A visual of our labeler program. The predicted trajectories from our baseline or AI algorithm can be manually corrected using this interface. It is designed for easy visualization of tracks and correction of crossovers.

The model parameters *v*, *σ_r_*, *σ_θ_*, and Ω are fit as population distributions from the real trajectories in our dataset. For each tracked, non-stationary sperm, we first smooth the raw centroid path with a centered rolling average (0.75 s window) and compute motion statistics from the smoothed path, since frame-to-frame centroid steps at our frame rate are comparable to detection noise and otherwise inflate heading and speed fluctuations. The chirality Ω and turning noise *σ_θ_* are estimated per sperm by fitting a two-Gaussian mixture, centered at *±*Ω, to the histogram of angular velocities, discarding the worst quartile of fits. Because the average path velocity (VAP) and the speed variability are derived statistics of the motion rather than direct readouts of *v* and *σ_r_*, they are biased estimates of these parameters (in our tests, VAP overestimated *v* by roughly 1.6–1.8*×*). We therefore built calibration curves by simulating trajectories at known values of *v* and *σ_r_*, measuring the same statistics on the output, and interpolating each real measurement back to an estimated true parameter value. Estimates pooled across all clips were trimmed to the 1st–99th percentile and fit with a normal distribution for Ω and lognormal distributions for *v*, *σ_r_*, and *σ_θ_*. The flagellar beating parameters (*ω*, *τ*) could not be recovered from our recordings. At 9 fps the Nyquist limit is 4.5 Hz, and simulated beat frequencies above this limit fold back to lower apparent beat-cross frequencies, so the mapping is not invertible. These parameters were instead fixed to constant values (*τ ≈* 1*/*3 s) with little spread.

To implement the equations and create the synthetic trajectories, we built a software package in Python using the NumPy library. The software allows for a user-defined number of sperm to exist in the video and their movement is calculated based on sampled parameters from the fitted probability distributions. Each trajectory is then computed for each individual synthetic sperm. To create a realistic video, the OpenCV package is used to draw the synthetic sperm on to each frame of the video. An ellipsoidal shape is drawn at each sperm’s given location, with the major axis facing towards the current movement direction. Additionally, randomness is added to the overall width and brightness of each sperm to imitate phenomena in real microscopy videos.

**Fig 3.**
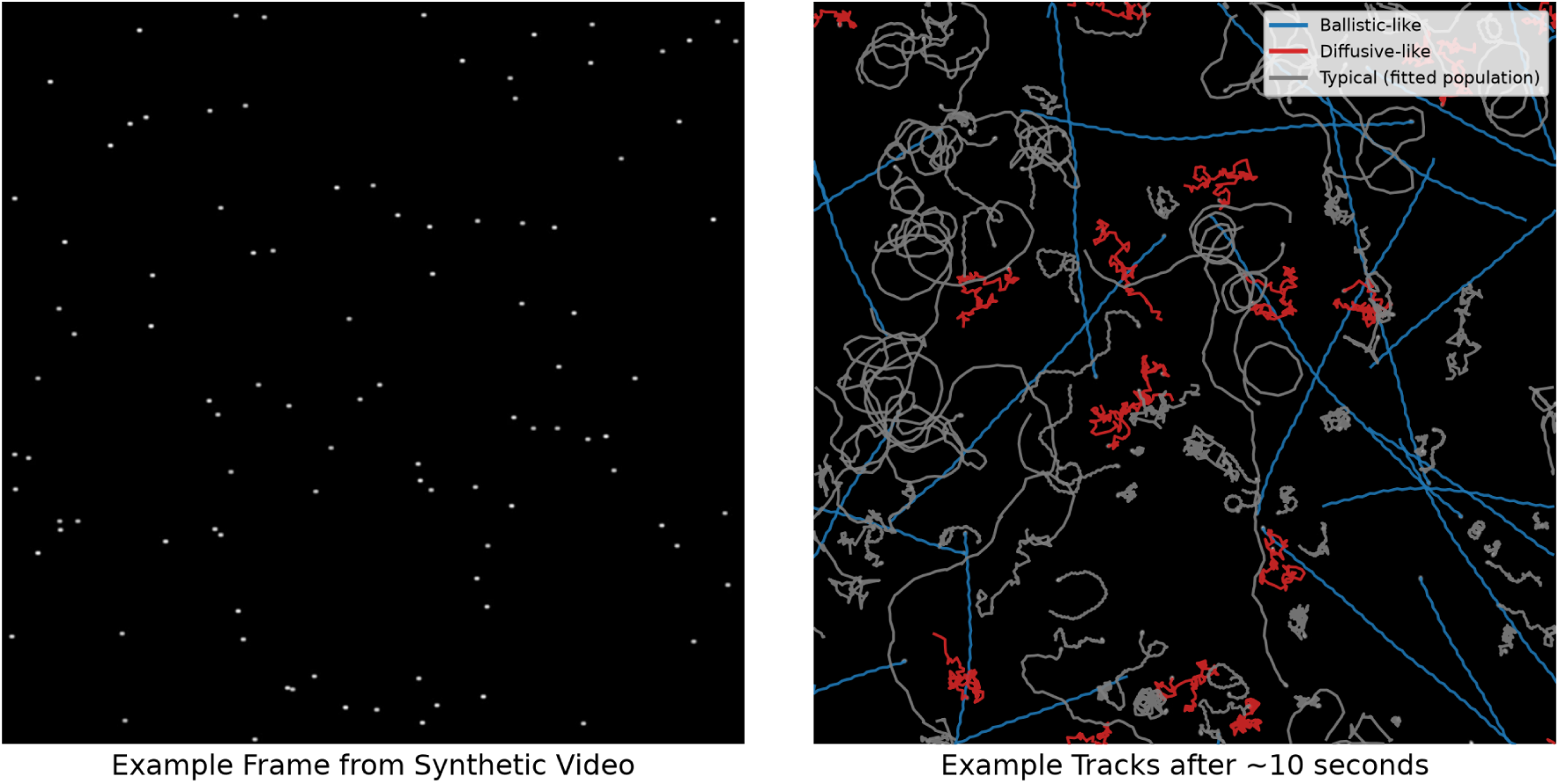
(Left) An example frame from a synthetically generated video. (Right) Example tracks of synthetic sperm after 10 seconds. The synthetic movement equations are fitted based on real labeled sperm data, and have the flexibility to represent many motions, ranging from “diffusive-like” to “ballistic-like” behavior.

This synthetic generation process allows for the creation of as many sperm microscopy videos as needed for the training pipeline. Additionally, because the number of sperm is user-defined, the crowdedness of the video can be directly controlled for effective training and evaluation.

### Augmenting Real Videos

In addition to the synthetic data, additional data can be generated from the real collected videos using augmentation techniques. Although it is only possible to initially label low-density videos in a reasonable time with the baseline tracker, high-density videos can be created by combining multiple low-density videos together. Since the videos in our dataset contain bright sperm objects on dark backgrounds, simply adding videos together frame-for-frame makes a new video of higher density that is also valid. Additionally, since the videos are square in resolution (1024 *×* 1024), the videos can be rotated and mirrored before addition to create additional variations of the content. A visualization of this process is shown in Fig. 4.

The .csv files for the individual videos can also be combined to maintain labeled ground truth data. A sperm ID renumbering process occurs to guarantee unique identities for all sperm in the combined video. Additionally, any video transformations must also be applied to the .csv data appropriately (e.g. Mirroring about the y-axis in the video would mirror the x-values about 512 in the .csv file). Overall, this process allows for realistic videos at higher densities with fully-labeled trajectories.

### Training the AI Model

With additional and higher density videos available, there is now sufficient training data to train the AI model. This work adopts a detector-tracker pipeline consisting of a YOLO-based object detector [27] followed by the ByteTrack multi-object tracking algorithm [28]. In this framework, the detector is first used to localize sperm heads independently in each frame, producing a set of bounding-box detections with confidence scores. These frame-level detections are then passed to ByteTrack, which associates detections across consecutive frames in order to assign persistent identities and recover full trajectories over time.

**Fig 4.**
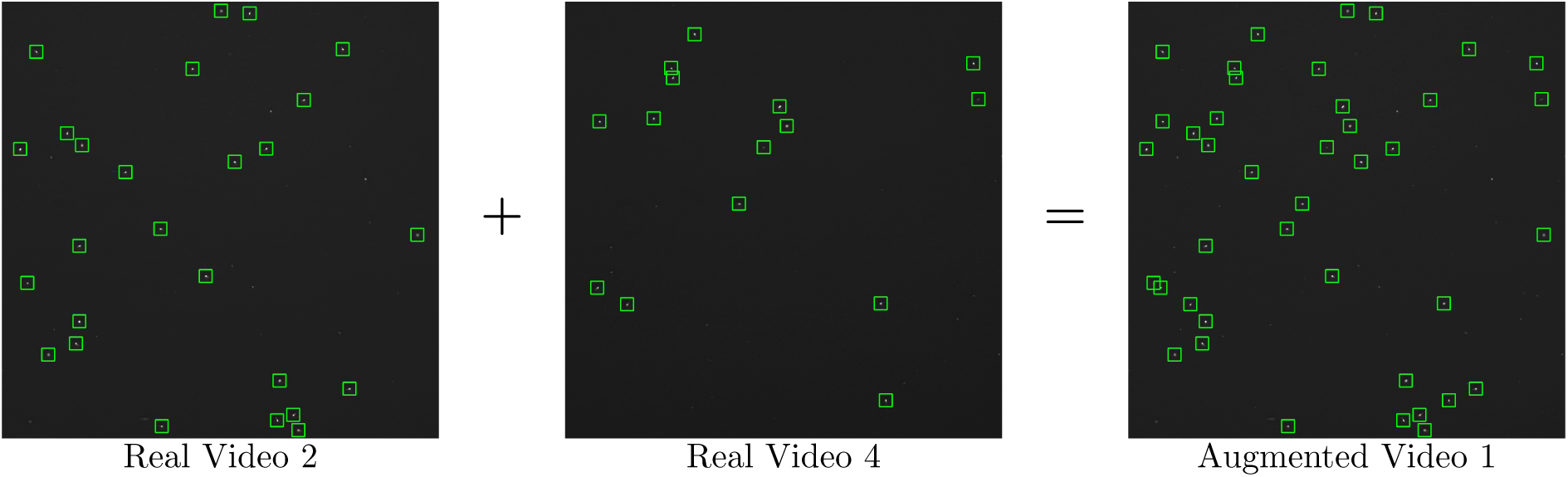
Example of an augmented video sequence (Right) generated by combining labeled Real Video 2 (Left) and Real Video 4 (Middle).

This detector-tracker decomposition is well suited to the sperm tracking problem. The detector is responsible for identifying sperm locations under challenging imaging conditions such as low contrast, variable brightness, and dense spatial packing, while the tracker is responsible for preserving temporal continuity as sperm move, cross paths, and occasionally disappear or reappear between frames. Separating detection from trajectory association makes it possible to study tracking quality directly and to evaluate how detection errors propagate into downstream trajectory-based measurements.

ByteTrack was selected because it is designed to improve identity association by using not only high-confidence detections, but also lower-confidence detections that would often be discarded by conventional tracking-by-detection pipelines [28]. This is especially relevant in crowded sperm microscopy videos, where a sperm cell may appear faint, partially obscured, or difficult to localize consistently in every frame. By retaining and associating weaker detections when appropriate, ByteTrack can reduce missed associations and improve track continuity relative to approaches that rely only on a strict confidence threshold.

The use of a YOLO-based detector is also consistent with recent sperm imaging studies that have applied one-stage object detectors to sperm head localization and have recovered motility patterns that remain comparable to those derived from ground truth trajectories [29, 30]. This supports the suitability of YOLO-style detectors as the front end of an automated sperm tracking pipeline.

### Training and Model Configuration

The YOLO detector was trained on ground-truth detection frames pooled from the labeled real videos, the synthetically generated videos, and the augmented real videos described above. Each labeled sperm centroid was converted to a fixed 30 *×* 30 pixel axis-aligned bounding box, rather than a box fit to the individual sperm’s apparent size or orientation, since the detector’s role in this pipeline is head localization rather than morphological characterization. All frames were treated as a single object class (*sperm*). Detector training and inference operate independently on individual frames. No temporal information is used during training, and video context is only introduced afterward by the tracking stage.

We fine-tuned a YOLO11x model [27], initialized from COCO-pretrained weights, at an input resolution of 1024 *×* 1024 pixels matching the native frame size of our recordings. We trained for 50 epochs with a batch size of 8 and an initial learning rate of 0.01 (SGD/AdamW, chosen automatically by Ultralytics), using the framework’s default augmentation settings (random horizontal flip, mosaic, and HSV color jitter). The training and validation sets were split at the video level rather than the frame level, so that every validation frame came from a video withheld from training entirely; this avoids the frame leakage that would result from a naive random split of highly correlated frames drawn from the same clip.

At inference time, frame-level detections above a confidence threshold of 0.40 were passed to ByteTrack [28] for identity association, using the tracker’s default configuration (first-stage association threshold 0.25, second-stage low-confidence threshold 0.10, a new-track threshold of 0.25, a track buffer of 30 frames, and an IoU association threshold of 0.80).

### Evaluation Metrics

With the newly trained YOLO+ByteTrack model, we verify its improved performance over the baseline tracking algorithm. We use the metrics defined from our previous work [20], but we briefly summarize them here for convenience.

The metrics come from multi-object tracking benchmarks for two major groups: natural image tracking [31] and cell tracking [32]. These metrics focus on different components of the tracking pipeline. Some metrics focus solely on the accuracy of the detection step (DET [33]), some focus on the accuracy of the linking/tracking step (IDF1 [34], LNK [33]), and other combine the detection and tracking accuracy together in a combined metric (MOTA [35], HOTA [36], TRA [33]). The metrics are computed differently based on how they weigh factors such as ID switches, missed detections, broken links, etc. One other important metric that is reported is track fractions (TF) which describes the average length of trajectory that is correctly identified by a predicted trajectory.

In addition to these previously reported metrics, we introduce one additional metric in this work that focuses on the models ability to resolve crossovers since this is a vital property of an effective assistive labeler. Crossovers are difficult to determine exactly, but likely candidates for a crossover can be estimated based on spatial and temporal proximity with some probability. To identify candidate crossover events, we examined every pair of sperm in the ground-truth trajectories and extracted proximity windows that indicated maximal contiguous runs of frames during which the pair’s centroid-to-centroid distance remained below a fixed threshold (25 pixels). Each window was scored using three geometric signals: whether the pair’s frame-to-frame displacement segments intersected at any point in the window (a true spatial crossing), whether sperm B’s position changed sides relative to sperm A’s direction of travel between the window’s entry and exit (a position swap, indicating a reversal of relative ordering without a literal path intersection), and the pair’s minimum distance within the window relative to the threshold. These were combined into a single likelihood score, *L ∈* [0, 1], defined as *L* = 0.6 + 0.4*p* when the segments intersect, *L* = 0.35 + 0.35*p* when only a position swap occurred, and *L* = 0.3*p* otherwise, where *p* = max(0, 1 *− d*_min_*/τ*) is the normalized proximity and *τ* is the 25 px threshold. These weights were chosen heuristically to impose a qualitative ranking rather than fit to labeled outcome data, but provide a relative confidence score. Tracking failures (identity switches or dropped detections) were then identified using a continuity-preserving match between ground truth and predicted trajectories. The number of possible crossover events and the number of failures in resolving the crossover event are reported in our results.

### The Data Engine Loop and Software Program

The result of the data labeling, synthetic and augmented data generation, and training steps described in the previous steps is an AI-based tracking model that can handle higher density video than the baseline tracker. With this AI tracker now available, trajectories for higher density video can be estimated with fewer corrections needed from a human labeler.

As shown, this data engine loop contains multiple steps and requires organizing multiple sources of data.

To simplify this process for ourselves and others, we have created a software program in PyQt [24] that implements an intuitive interface for completing all the steps of the data loop. Functionality and features include:

- Initializing the labeler program with predicted tracks from a specific tracking algorithm and weights.
- Automatic synthetic parameter fitting and streamlined controls for new synthetic and augmented video creation.
- A video registration menu to assign videos from real, synthetic, and augmented sources to training and validation datasets.
- A YOLO+ByteTrack training status bar for realtime monitoring.
- Built-in evaluation and visualization functionality for post-training analysis.

This functionality makes the data engine loop easy to follow. Our full program is documented and provided in our GitHub repository at https://github.com/CAS-ReproLab/Sperm_Object_Tracking/.

## Results

With the data engine pipeline established, we now report the evaluation metrics on the baseline and YOLO+Bytetrack approaches for videos of various sperm densities.

### Evaluating the Trained AI Model

For purposes of the provided intuition to the results, we binned the videos based on density and labeled these videos as low density if they contained less than 80 sperm trajectories, medium density if they contained 80 to 240 sperm trajectories, and high density if they contained more than 240 sperm trajectories. These groups were chosen based on densities roughly found at different dilution levels in the microscopy videos.

The first iteration of the data engine loop took the 5 initial low-density labeled videos, with corresponding synthetic and augmented videos generated at medium density, and generated a training/validation split. The training split consisted of 4 real low-density videos, 3 synthetic medium-density videos, and 3 augmented medium-density videos. The validation split consisted of 1 real low-density video, 2 synthetic medium-density videos, 2 augmented medium-density videos, 1 synthetic high-density video, and 1 augmented high-density video. this represents a total of 2132 sperm in the training set and 2903 sperm in the validation set.

The YOLO+ByteTrack approach is compared to the baseline Trackpy method. Both trackers were evaluated on the validation set using the benchmark metrics. The results of these findings are shown in Table 1 and further visualized in Figure 5.

**Table 1.** Evaluation of results on sperm trajectory data across low, medium, and high density conditions (filtered to include only moving sperm), comparing the Trackpy baseline against the trained YOLO+ByteTrack model.

| **Density - Type** | **Tracker** | **DET** | **TRA** | **LNK** | **TF** | **MOTA** | **IDF1** | **HOTA** |
|---|---|---|---|---|---|---|---|---|
| Low - Real | Trackpy | 0.973 | 0.975 | 0.983 | 0.866 | 0.841 | 0.814 | 0.839 |
|  | YOLO+ByteTrack | **0.986** | **0.986** | **0.984** | **0.902** | **0.941** | **0.951** | **0.949** |
| Medium - Synthetic | Trackpy | 0.949 | 0.947 | 0.934 | 0.735 | 0.949 | 0.704 | 0.740 |
|  | YOLO+ByteTrack | **0.991** | **0.990** | **0.986** | **0.877** | **0.984** | **0.878** | **0.910** |
| Medium - Augmented | Trackpy | 0.952 | 0.954 | **0.969** | 0.814 | 0.715 | 0.711 | 0.742 |
|  | YOLO+ByteTrack | **0.970** | **0.970** | 0.964 | **0.836** | **0.936** | **0.941** | **0.935** |
| High - Synthetic | Trackpy | 0.924 | 0.921 | 0.897 | 0.624 | 0.910 | 0.597 | 0.645 |
|  | YOLO+ByteTrack | **0.979** | **0.978** | **0.972** | **0.856** | **0.972** | **0.870** | **0.892** |
| High - Augmented | Trackpy | 0.831 | 0.823 | 0.790 | 0.388 | 0.634 | 0.445 | 0.487 |
|  | YOLO+ByteTrack | **0.846** | **0.844** | **0.829** | **0.607** | **0.778** | **0.758** | **0.751** |

### Capability in Resolving Crossovers

The tracking metrics presented are helpful indicators of the AI model’s performance, but do not directly correlate to usefulness in the labeling process. The most common error that needs correction during the labeling process is the loss or switch of sperm IDs when two sperms crossover each other the video. Thus, we compare the the initial Trackpy tracking algorithm to the trained YOLO+ByteTrack algorithm on their ability to resolve the possible crossovers in the labeled data. The results are shown in Table 2. As is shown, the AI model does significantly better than the Trackpy model across all densities. Specifically, the trained AI model had the crossover failure rate on medium-density data going from 44.8% to 6.64% on medium density videos, and from 74.0% to 46.68% on high density videos. This eliminating of crossovers significantly increases the speed of the labeling process for sperm trajectories.

**Fig 5.**
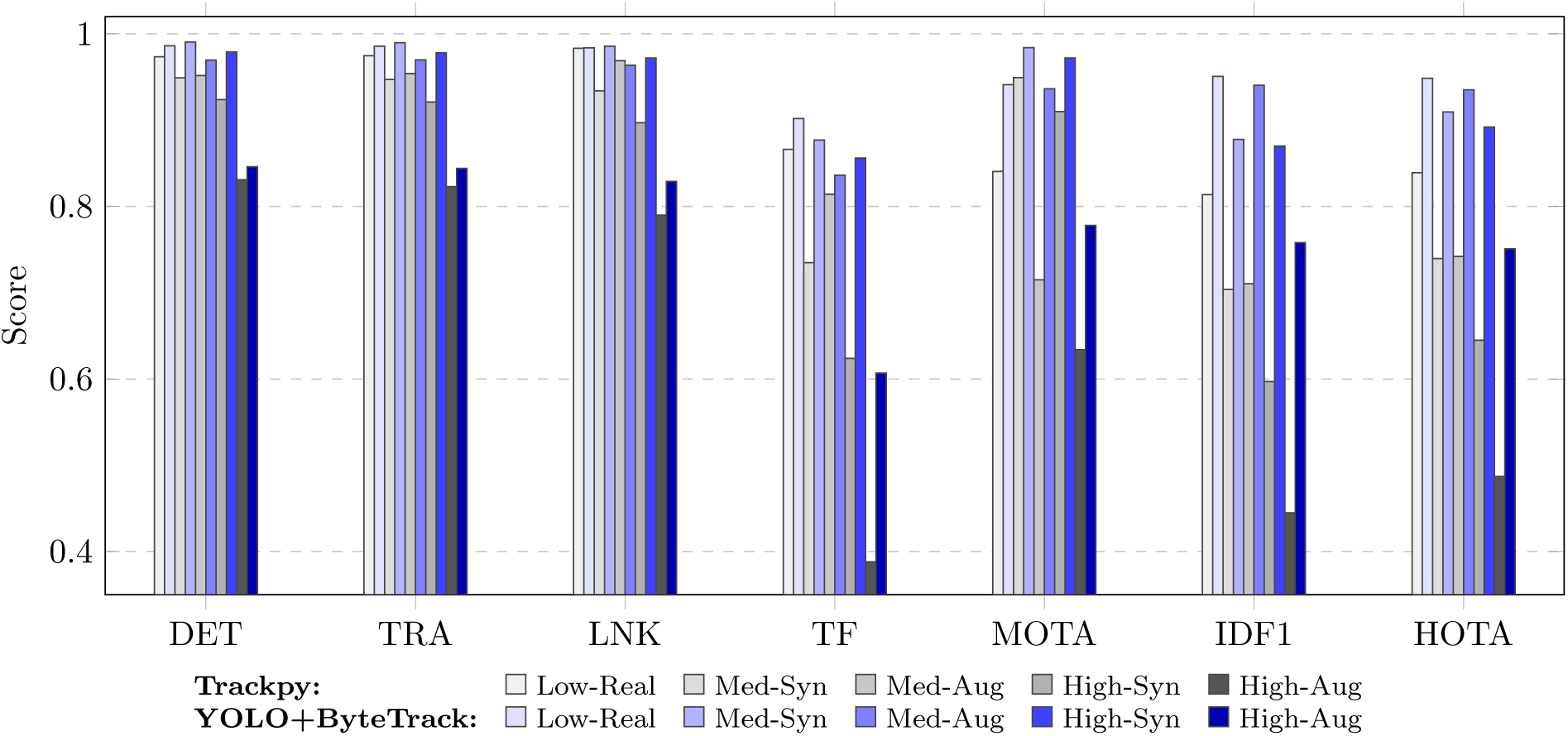
Evaluation on filtered sperm trajectory data, comparing Trackpy (gray) against YOLO+ByteTrack (blue). Within each metric, bars are paired by condition (Trackpy left, YOLO+ByteTrack right) in the order Low-Real, Medium-Synthetic (Med-Syn), Medium-Augmented (Med-Aug), High-Synthetic (High-Syn), and High-Augmented (High-Aug).

**Table 2.** Comparison of the Trackpy baseline against the trained YOLO+ByteTrack model in capability in correctly tracking sperm through crossover events.

| **Density - Type** | **Tracker** | Crossover Events | Tracking Failures | Failure Rate |
|---|---|---|---|---|
| Low - Real | Trackpy | 68 | 24 | 0.3529 |
|  | YOLO+ByteTrack | 68 | **1** | 0.0147 |
| Medium - Synthetic | Trackpy | 1535 | 832 | 0.5420 |
|  | YOLO+ByteTrack | 1535 | **196** | 0.1277 |
| Medium - Augmented | Trackpy | 241 | 108 | 0.4481 |
|  | YOLO+ByteTrack | 241 | **16** | 0.0664 |
| High - Synthetic | Trackpy | 3065 | 1957 | 0.6385 |
|  | YOLO+ByteTrack | 3065 | **583** | 0.1902 |
| High - Augmented | Trackpy | 4659 | 3447 | 0.7399 |
|  | YOLO+ByteTrack | 4659 | **2175** | 0.4668 |

### Final Outputs: Labeled Medium and High Density Data

With the AI-based assisted-labeler, higher density videos are able to be labeled in significantly shorter times. Three additional medium density videos were able to labeled with an average of user hand-correction time of around 30 minutes, and 1 additional high density video was labeled with a total correction time of around 4 hours. In comparison, based on number of possible crossover events and Trackpy failure rates, hand-correction times would likely be 3-5 hours for medium-density data and 6-12 hours for high-density data. While additional user studies would be needed to validate these estimates, this represents a significant speed-up in the labeling time process.

With this paper, we included the original 5 low-density videos, the 3 newly labeled medium-density video, and 1 fully labeled high-density video. A visual example of the fully labeled high-density data is shown in Figure 6.

**Fig 6.**
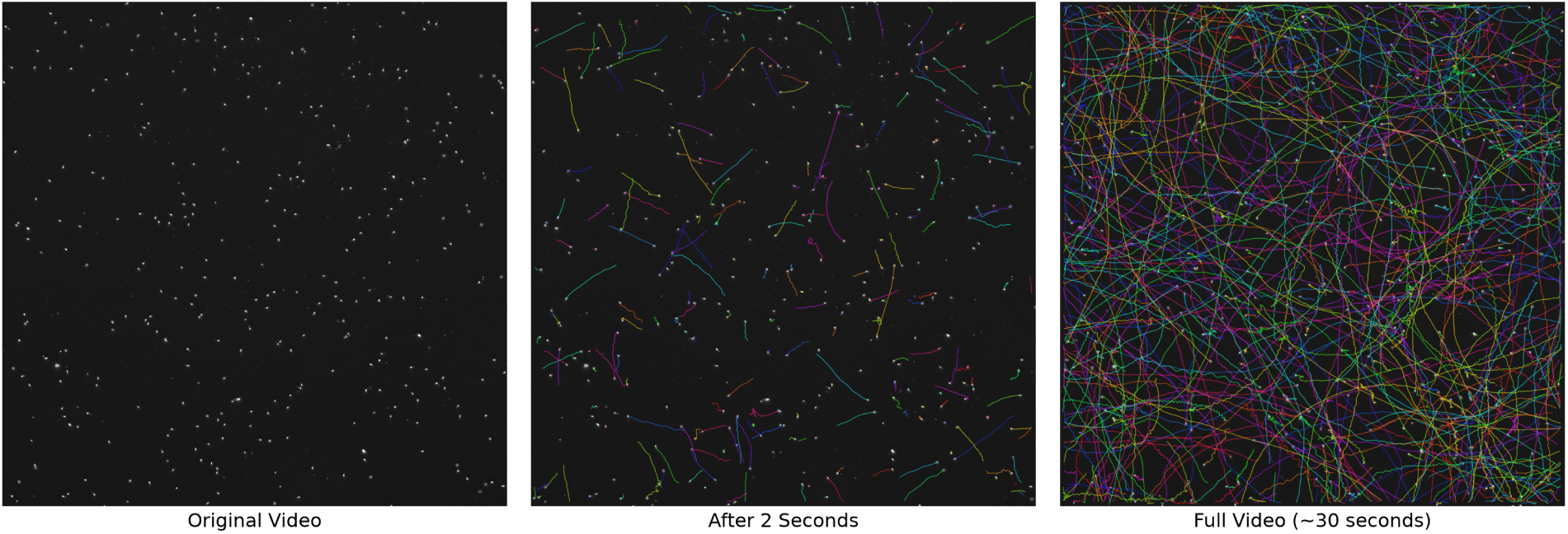
A visual of a fully labeled high-density video. For traditional sperm tracking datasets and algorithms, it is common to only capture 2 seconds of video to limit the number of cross overs. But with our new approach, we can now visualize and analyze the trajectories at densities and lengths of time not previously possible.

### Ablation: Another Iteration of the Data Loop

Using the Loop-1 tracker for assisted labeling, we labeled three additional real videos at medium density. We then investigated whether a second round of training on these new data would improve the model’s usefulness for labeling even higher-density videos.

Loop 2 of the data engine started from the complete Loop-1 dataset (all training and validation videos). To the training set we added the two newly labeled real medium-density videos, along with one synthetic and one augmented high-density video. We compared three training strategies, which differ in how the model was initialized and which data it saw. Model A was trained from scratch on all data. Model B was initialized with the Loop-1 weights and fine-tuned on all data. Model C was initialized with the Loop-1 weights and fine-tuned only on the new data (i.e. the real medium-density videos and the generated high-density videos).

The Loop-2 YOLO+ByteTrack approach and its variations are compared to the baseline Trackpy method and Loop-1 tracker. The results of these evaluations on the baseline metrics are shown in are shown in Table 3. Additionally, the crossover analysis for this stage is given in 4.

As can be seen, even with the additional training data, the Loop 2 model did similarly or worse compared to the original Loop 1 model. This is likely due to a limitation in the tracking pipeline algorithms. The common YOLO+Bytetrack setup separates the detection and linking phases. Objects are found in each frame and then followed from frame to frame. In these setups, the YOLO model is the detection step and learned component from the AI. ByteTrack is the linking step and is not learned, but instead relies on motion priors and Kalman filters. This limits the models ability to follow the hardest crossover motions, even if detection is happening perfectly. Future AI models that can learn the motion priors are likely to improve with additional data loop iterations.

## Discussion

Computer-aided sperm analysis (CASA) has evolved alongside advances in microscopy and computing, from manual trajectory reconstruction to automated image analysis and modern machine learning approaches [37]. Despite these advances, most CASA systems remain optimized for short recordings of diluted samples, partly because tracking algorithms struggle to preserve sperm identities when trajectories intersect [38]. Here, we build on our previous study that established effective tracking metrics for sperm microscopy videos [20]. We use Trackpy [23], a classical particle-tracking package, as the initial tracking model for the low-density videos. We developed our own software for labeling, synthetic video generation, and augmentation of real videos. We then trained a YOLO-based object detection model [27] on the generated dataset and paired it with ByteTrack [28], a tracking-by-detection method that associates detections across frames, including low-confidence ones. This tracker was then used for improved assistive labeling of microscopy videos with increasingly high sperm densities. We evaluated the AI tracker directly with multiple metrics that were proposed in our previous work [20]. Compared with conventional particle tracking, our approach reduced crossover failure rates from 44.8% to 6.64% in medium-density augmented videos and from 74.0% to 46.68% in high-density augmented videos, substantially reducing the effort required to construct labeled trajectory datasets.

**Table 3.**
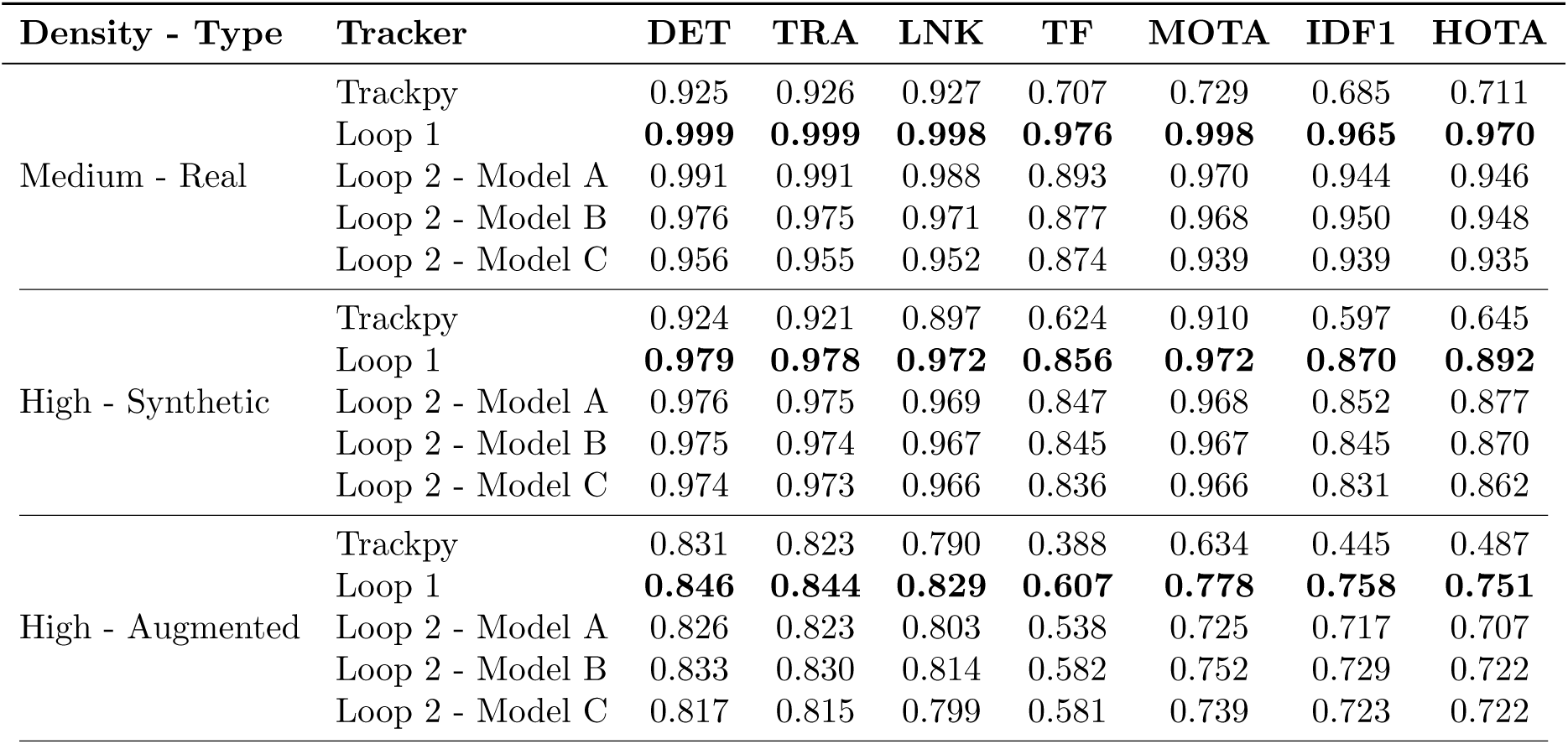
Evaluation of results on filtered sperm trajectory data across medium and high-density conditions, comparing the Trackpy baseline and Loop-1 models against the trained Loop-2 models.

Recent machine learning approaches to semen analysis have focused primarily on improving sperm segmentation and detection using neural networks [39]. However, linking detected sperm into accurate trajectories still remains a major challenge. Although AI-based methods can outperform classical algorithms, their development depends on accurately annotated training data, which are particularly difficult to obtain from high density sperm microscopy videos characteristic of semen analysis. Our approach differs from other recent methods by addressing this data bottleneck. We aimed to develop a system of model training that is somewhat agnostic to the models used, rather than simply provide pre-trained models. Modern high-resolution digital microscopy, computational resources, and open-source software make it possible to generate large synthetic and augmented datasets, train neural detectors, and iteratively improve tracking through human-assisted correction. The resulting open datasets and software provide a foundation for developing and benchmarking more advanced tracking methods. Nevertheless, our second training iteration did not improve performance, suggesting that further progress may require learned motion models rather than improvements in detection alone.

In the 1950s, Austin and Chang independently observed that a period of residency in the reproductive tract is necessary for mammalian sperm to gain fertility competence [40, 41]. During this residency, sperm undergo a series of biochemical changes known collectively as ‘capacitation’, which include acrosomal exocytosis, remodeling of molecular surface-binding properties, changes in metabolic flux patterns, and shifts in kinematics [42]. The rate and degree of capacitation varies from cell to cell, and significant heterogeneity among sperm subpopulations has been observed across mammalian species [43–46].

A few promising studies point to the validity of connecting sperm kinematics and other biochemical changes to random search behavior. Zaferani and colleagues elegantly showed that bovine sperm exhibit ‘biphasic chemokinesis’ in response to capacitative signals in vitro [47]. They then followed that work with a recent study that used single cell trapping in a microfluidics device to facilitate trajectory reconstruction over more than ten minutes, revealing long timescale phase transitions between ‘circling and wandering’ behaviors that were predicted to optimize first coverage time in complex search environments [19]. These findings are consistent with our recent biophysical modeling study, wherein we showed that phase transitions from high to low persistence behavior maximize area coverage during random search [18]. Additionally, Giojalas and Eisenbach explored sperm subpopulation-scale variation in acrosome reaction rate confers an adaptive statistical ‘anticipation’ of the time or distance to a fertilizable egg [48]. Despite the emerging evidence, sperm heterogeneity and long timescale behaviors have not been widely incorporated into CASA analysis or male fertility assessment.

**Table 4.** Comparison of the baseline, Loop-1, and Loop-2 models in capability in correctly tracking sperm through crossover events.

| **Density - Type** | **Tracker** | Crossover Events | Tracking Failures | Failure Rate |
|---|---|---|---|---|
| Medium - Real | Trackpy | 665 | 363 | 0.5459 |
|  | Loop 1 | 665 | **41** | 0.0617 |
|  | Loop 2 - Model A | 665 | 61 | 0.0917 |
|  | Loop 2 - Model B | 665 | 88 | 0.1323 |
|  | Loop 2 - Model C | 665 | 101 | 0.1519 |
| High - Synthetic | Trackpy | 3065 | 1957 | 0.6385 |
|  | Loop 1 | 3065 | **583** | 0.1902 |
|  | Loop 2 - Model A | 3065 | 682 | 0.2225 |
|  | Loop 2 - Model B | 3065 | 726 | 0.2369 |
|  | Loop 2 - Model C | 3065 | 736 | 0.2401 |
| High - Augmented | Trackpy | 4659 | 3447 | 0.7399 |
|  | Loop 1 | 4659 | 2175 | 0.4668 |
|  | Loop 2 - Model A | 4659 | **2141** | 0.4595 |
|  | Loop 2 - Model B | 4659 | 2263 | 0.4857 |
|  | Loop 2 - Model C | 4659 | 2241 | 0.4810 |

Together, the emerging evidence points to a new frontier in the biology of male fertility, wherein sperm physiology and large numbers facilitate the solution to a complex random search problem, which predicts that target encounter probabilities depend on trajectory persistence, reorientation frequency, and environmental geometry, rather than speed alone [17]. Progress will require practical technologies that support long observation periods for sperm movement, along with new kinematic measures that link observed movement patterns to search behaviors.

By enabling longer observations at higher sperm densities, our approach could expand semen analysis beyond conventional short-term kinematic measurements. Longer trajectories would permit measurements of directional persistence, behavioral transitions, and spatial exploration, while higher-density recordings could reveal sperm–sperm interactions and population-level organization that are obscured by sample dilution. These measurements could help distinguish sperm populations with similar conventional motility parameters, but different transport and search behaviors. Although the present system remains limited by imaging conditions, temporal resolution, and residual tracking errors, and the relationship between these measurements and fertility outcomes remains to be established, the approach described here offers a path toward more physiologically informative semen analysis, connecting sperm motility to the biological processes required for successful fertilization.

### Limitations

The data generation process, both for synthetic and augmented videos, has inherit limitations for what it can represent. First, in the synthetic data, there is currently no mechanism for including debris or other material in the generated images. These distractor objects are commonly present in real microscopy videos and could be included to increase the realism of the synthetic data. Second, the data generation process is currently defined for negative phase contrast microscopy. Additional work would be needed to adjust the system for positive phase contrast microscopy, which often includes a grey background and “halo” effects around the sperm. Though not explored in this work, additional image processing techniques should allow for both the synthetic and augmented generation pipelines to work for positive phase contrast videos.

We also acknowledge a limitation in the kinds of motion the synthetic and augmented data generation process can represent. Those motions are either sampled from our assumed synthetic trajectory equations, or are literal copies from lower density data. Thus the motion statistics in the generated videos will not be as varied as true high density data. However, we note the purpose of this training data is not to capture broader motion statistics, but to simply create more crossovers as would be seen in higher density data. The AI model needs more visual examples of what sperm overlapping looks like in the video feed. Since resolving a crossover only requires a few frames before and after the event, simple motion models are generally sufficient given proper detection of the individual sperm.

Lastly, as mentioned in the Ablation section, the motion models ByteTrack uses cannot be adjusted and fail in the most difficult motion cases (e.g. cross-overs between two hyperactive sperm). For this problem to be deeply solved at the high densities, there needs to be a more effective mechanism for learning “motion” rather than leaning on simple motion models. Bytetrack ultimately does tracking by redetection, which can’t resolve these difficult cases with the limited visual signal provided by microscopy videos. The AI community will need to create tracking models that move beyond detect-link framework to to fully resolve this problem.

## Conclusion

This work provides five resources, all of which will be made publicly available: (i) video data collected across a range of sperm population densities; (ii) an assisted-labeling program designed specifically for sperm microscopy videos; (iii) corrected trajectory datasets for sperm videos at low, medium, and high densities; (iv) AI models for sperm tracking, with pretrained weights; and (v) a comprehensive software package that incorporates data labeling, synthetic and augmented video generation, AI training, and evaluation for sperm microscopy data. These are all included in our GitHub repository, along with documentation at https://github.com/CAS-ReproLab/Sperm_Object_Tracking/.

